# Early survival in Atlantic salmon is associated with parental genotypes at loci linked to timing of maturation

**DOI:** 10.1101/2024.01.08.574625

**Authors:** Tutku Aykanat, Darryl McLennan, Neil B. Metcalfe, Jenni M. Prokkola

## Abstract

Large effects loci often contain genes with critical developmental functions with potentially broad effects across life-stages. However, the life-stage-specific fitness consequences are rarely explored. In Atlantic salmon, variation in two large-effect loci, *six6* and *vgll3*, is linked to age at maturity, and several physiological and behavioural traits in early life. By genotyping the progeny of wild Atlantic salmon that were planted into natural streams with nutrient manipulations, we tested if genetic variation in these loci is associated with survival in early life. We found that higher early life survival was linked to the genotype associated with late maturation in the *vgll3*, but with early maturation in the *six6* locus. These effects were significant in high-nutrient, but not in in low-nutrient streams. The differences in early survival were not explained by additive genetic effects in the offspring generation, but by maternal genotypes in the *six6* locus, and by both parents’ genotypes in the *vgll3* locus. Our results suggest that indirect genetic effects by large-effect loci can be significant determinants of offspring fitness. This study demonstrates an intriguing case of how large-effect loci can exhibit complex fitness associations across life stages in the wild and indicates that predicting evolutionary dynamics is difficult.

## Introduction

Adaptation in the wild is a complex process, composed of interactions between many mechanisms, such as context-dependency of fitness across time and space. Understanding the genetic architecture of fitness has been one of the key aims of evolutionary quantitative genetics. Classical theory suggests that mutations with large adaptive effects may hinder fine scale adaptation (Orr 2000, 2005). Yet, several quantitative trait loci (QTLs) have been identified to be associated with fitness in natural systems (Bradshaw and Schemske 2003; Colosimo et al. 2005; Johnston et al. 2013; Tishkoff et al. 2007), indicating that adaptive genetic variation via large effect loci can persist. Orr (2000), on the other hand, suggests that genetic correlations between fitness components should be selected against. However, Yeaman and Whitlock (2011) predict that under certain migration-selection scenarios, adaptive loci will be selected for to cluster tightly within a genomic region. As such, adaptive genomic regions with large phenotypic effects may contain multiple loci associated with different fitness components, resulting in high genetic correlations among them. How such genetic architecture is tied to physiological processes remains poorly understood. For example, a genomic region with a large phenotypic effect may be centrally located within cellular and developmental networks, and the genetic variation may in turn be linked to broad changes in body plan and/or physiology (Chan et al. 2010; Colosimo et al. 2005; Jimenez-Gomez et al. 2010; Snoek et al. 2012; Weith et al. 2023). Such broad expression of phenotypes is likely to alter fitness via different pathways, perhaps across different tissues or life stages. Yet, little is known of the fitness landscapes of large effect loci across life stages in the wild. Determining the direction of genetic correlations between fitness components and their potential costs across different environmental conditions is crucial to a better understanding of how genetic variation is maintained in populations, and of evolutionary responses to environmental change.

The Atlantic salmon has been emerging as a promising wild model to explore the multitude of fitness effects associated with large effect loci. Wild salmon populations are adapted to their local environments, so that even those in close proximity and lacking barriers to migration can exhibit genetic and phenotypic differences (Taylor 1991; Garcia de Leaniz et al. 2007). At the genetic level, two genomic regions, so-called *vgll3* and *six6* (named after the most predominant genes within the loci, *vestigial-like family member 3* and *SIX homeobox 6*, respectively) exhibit substantial signatures of adaptation within and among populations (Barson et al. 2015; Pritchard et al. 2018). These genomic regions harbour genes encoding transcription (co)factors, and are expressed as early as embryonic development (Halperin et al. 2013; Kurko et al. 2020; Lee et al. 2012; Moustakas-Verho et al. 2020). Genetic variation in the *vgll3* underpins key life-history traits in Atlantic salmon, such as the age at maturity in wild individuals (Ayllon et al. 2015; Barson et al. 2015) and in captivity (Asheim et al. 2023), the probability of males becoming precociously mature prior to sea migration (Debes et al. 2021), and iteroparity (Aykanat et al. 2019). *Vgll3* is also associated with physiological and behavioural traits in juvenile fish, such as lipid profiles (House et al. 2021), aggression (Bangura et al. 2022), and (sex-specific) smolt migration (Niemela et al. 2022), suggesting that this single QTL can have broad effects across multiple life stages. In addition, the genetic variation in these loci is associated with metabolic phenotypes, whereby juvenile salmon with the *vgll3* early genotype exhibit a higher maximum metabolic rate and aerobic scope, with the former trait also affected by *six6* loci via epistasis (Prokkola et al. 2022). These genomic regions also show physiological epistasis via the maximal activities of mitochondrial and anaerobic metabolic enzymes in the heart and intestine (Prokkola et al. 2024). Variation in *six6*, in turn, has been associated with stomach content volume during ocean feeding (Aykanat et al. 2020). Similar to *vgll3,* the allele frequencies of the *six6* region are highly correlated with the average age at maturity in wild populations (Barson et al. 2015), and explain variation in age-at-maturity of a domesticated aquaculture strain of Atlantic salmon (Sinclair-Waters et al. 2020), as well as in Pacific salmonids (Waters et al. 2021). Taken together, these studies suggest that genetic variation in these two loci may be linked to differences in resource acquisition and allocation in juvenile salmon, with cascading effects on life-history strategies.

The early life stages of Atlantic salmon are characterized by high rates of mortality, in large part due to intense density-dependent resource competition (Einum et al. 2006; Grossman and Simon 2019). The availability of resources such as food and territorial refuge may strengthen or weaken this competition. The likelihood of an individual surviving to maturation may depend on what resources are available and how capable that individual is at competing for such resources. Considering the associations of *vgll3* and *six6* with resource acquisition and allocation, we predict that these loci interact with variation in stream nutrient levels to predict survival rates of juvenile salmon. However, directional predictions on how freshwater conditions (i.e., nutrient availability) should lead to adaptive differences in the maturation age of Atlantic salmon (i.e., the orientation and strength of selection) are challenging to formulate. For example, favourable growth opportunities and increased freshwater growth could induce early maturation (Salminen 1997), but also reduce the age when salmon juveniles migrate to sea, which imposes an opposite effect on maturation timing (Marschall et al. 1998). Such physiological complexities, coupled with context-dependent adaptive responses, make it difficult to formulate *a priori* predictions on the gene-phenotype relationship in relation to the adaptive significance of maturation timing (Barrett et al. 2009; Reznick 2016).

In this study we test whether the *vgll3* and *six6* loci are linked to early life survival in Atlantic salmon. We used data from a field experiment in which fertilized eggs from the same wild-origin families were divided among 10 experimental streams, all of which had become increasingly oligotrophic in recent years as a result of land management practices and anthropogenic barriers to the migration of anadromous fish (e.g., Bernthal et al. 2022). Five of these streams were given an experimental supplement to restore nutrient that imitates conditions of natural spawning levels prior to the river became inaccessible to migrating salmon (Auer et al. 2018). A previous study using the same field system showed that family-level juvenile survival was influenced by this manipulation of stream nutrient availability in addition to family-level maximum metabolic rate and egg mass (Auer et al. 2018). In this current study, we test for associations between juvenile survival and genetic variation in the two life-history loci, *vgll3* and *six6*. More specifically, we first tested whether differences in survival are due to family-wise genetic effects, i.e., associated with family-level allele frequencies of *vgll3* and *six6*, and second, we tested for direct additive effects of the loci, and parental effects, as occurs when the outcome on offspring survival depends on which parent passed on the allele.

## Materials and Methods

The full field protocol is described by Auer et al. (2018). In brief, wild adult Atlantic salmon were trapped on their upstream spawning migration as part of fishery management operations in the River Conon catchment in northern Scotland in late autumn 2015 and used to generate 30 full-sib crosses (different parental fish were used for each full-family) by using *in vitro* fertilization. Genotype data were lost for one family, and so 29 families were used in all analyses in the present study. All parental fish were selected to be fish that had spent one winter at sea (the most common marine life history phenotype in this catchment, subsequently confirmed by scale reading) to avoid major differences in egg (and hence initial offspring) size, which could affect among-family differences in early life survival. Egg mass was estimated for all families using a subset of eggs. Parents were weighed (to nearest 10 g) and measured for fork length (to nearest 5 mm), and a fin clip was taken for DNA analyses, i.e., for the parental assignment of each of the recaptured offspring (see below). Condition factor was calculated using a modified version of Fulton’s condition factor, CF = 100 x Weight/(Length)^m^, where *m* is the slope coefficient of the linear regression between log(weight) (response variable) and log(length), as the covariate (Bolger and Connolly 1989). This formula corrects for the residual relationship between length (or weight) and condition factor, hence condition factor can be modelled independently from the length of individuals.

Eggs were initially reared as family-specific groups and were overwintered in a local hatchery until the eyed stage of development, when a subsample of eggs from each of the full-sib families were transferred to the aquarium facilities at the University of Glasgow to be reared under standardized conditions. Hatchery mortality of eggs and subsequently of fry in the aquarium was negligible (<5%). Meanwhile, another subsample of 100 eggs from each of the same families were planted at the same density in each of 10 streams (in the same River Conon catchment in which the parent fish had been caught) in February/early March 2016, thus equalling 3,000 eggs per stream. There was no natural spawning of Atlantic salmon in these streams due to the presence of downstream dams that prevented the upstream migration of spawning fish. All ten streams were oligotrophic at the start of the experiment. Half of these streams were then experimentally supplemented with nutrient pellets at the time of egg planting, thereby simulating the decay of parental carcasses (referred to as the high nutrient streams, Auer et al. 2018), whereas nutrient levels in the other five streams were unaltered (referred to as the low nutrient streams). Previous ecological analyses of data from this experiment have shown that these nutrient additions led to significant increases in macroinvertebrate densities, and consequently faster growth of the young salmon (details in Auer et al. 2018; McLennan et al. 2019).

The maximum metabolic rate (MMR) and standard metabolic rate (SMR) of 10 individuals from each family were measured using continuous flow-through respirometry in late June 2016, using the offspring that had been reared under common garden conditions (Auer et al. 2018) following protocols from (Auer et al. 2015). Standardized triple-pass electrofishing surveys were conducted in each of the experimental streams in July 2016, when the fish were approximately three months post-first-feeding (Auer et al. 2018), to calculate-family level rates of growth and survival. A small sample of adipose fin tissue was taken from the recaptured individuals to allow parental (and thus family) assignment (Auer et al. 2018). In the present study, we used these same DNA samples of the parental fish and of the offspring that survived in the streams to genotype individuals at the *vgll3* and *six6* loci, using kompetitive allele specific PCR (KASP) method, as described in Prokkola et al. (2022). Throughout the paper, we refer to the alleles associated with early and late maturation as E and L, respectively. Consequently, EE and LL will be used for fish that are homozygous for E and L alleles and EL for heterozygotes, respectively, for both loci (so that the possible genotypes are *vgll3^EE^, vgll3^LL^, vgll3^EL^*, and *six6^EE^, six6^EL^, six6^LL^*). Out of 1246 juveniles that were collected and genotyped, the genotyping success was 96.3% and 97.5% for *six6* and *vgll3* genotypes, respectively. Three individuals were scored with improbable genotypes (one for *six6*, and two for *vgll3* loci) which did not conform to the Mendelian expectations based on parental genotypes. The genotype of one of these individuals, which could not be inferred, was excluded, while the genotypes of two individuals were corrected to the genotype value that was invariable within the full-sib family. Across the 58 parents that were used to generate the 29 families, the gene frequencies were unbalanced towards the early genotypes (0.85 and 0.63 for *vgll3*^E^ and *six6*^E^, respectively, see Supplementary Table 1), likely due to selecting parents that matured after one year at sea, and of the bias towards fish maturing after one sea winter in this population. In particular, for the *vgll3* locus, there were no *vgll3^LL^* individuals, and only one family (out of 29) that could generate *vgll3^LL^* offspring (i.e., family 9 in Supplementary Table 1).

### Family-wise early life survival in relation to life-history genotype

We performed a generalized linear mixed model to quantify survival among full-sib families within the high and low nutrient streams in relation to their genotype. We measured local survival as the number of individuals recaptured from the 100 eggs per family that were planted in each stream. While emigration is possible, juvenile salmon are predominantly sedentary in their first year of life, with limited movement and a high rate of mortality (Saunders and Gee 1964), suggesting that the local recapture rate can be mostly attributable to survival rather than emigration. Models were run using the *glmmTMB* package in R version 4.2.2 (Brooks et al. 2022; R Core Team 2021) and fitted using a negative binomial error structure with quadratic parameterization of mean in relation to variance (the *negbin2* family structure in the *glmmTMB* function).

The full model structure is as follows:

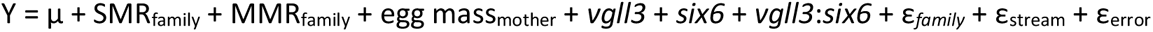

Whereby Y is the number of surviving individuals per family and stream, μ is the intercept, and *vgll3* and *six6* are the family-wise expected genotype frequency in these genomic regions, and *vgll3:six6* indicating the interaction term between *vgll3* and *six6*. In addition, the model included SMRfamily, MMRfamily, and egg massmother which are family-wise mean values for maximum metabolic rate (MMR) and standard metabolic rate (SMR), and mean egg mass per mother, respectively. These physiological indices were added as covariates, since Auer et al. (2018) showed that family-wise local survival was positively related to family-wise MMR and egg mass in the low nutrient streams, and marginally positively related to SMR in the high nutrient streams.

Family-wise expected genotype frequencies were modelled continuously, and calculated based on parental genotypes and segregation of alleles according to the Mendelian inheritance principles. For example, in a family where the parents’ genotypes are EE (0) and EL (0.5), the family-wise expected genotype frequency is 0.25, with 50% probability of offspring to be EE and 50% probability to be EL (see Supplementary Table 1 for parental genotype information). Family and stream were included in the model as random terms to account for non-independence among individuals of a given family or in each stream.

We performed a semi-automated model evaluation, using the *dredge* function in the *MuMin* package (version 1.47.1, Barton 2022) in R, by which we evaluated all possible fixed effect structures reduced from the full model, and ranked their fit according to the AICc value (a version of Akaike information criterion that corrects for small sample size), in which models within ΔAICc < 2 to the best model considered equally parsimonious. In a few instances, where convergence have not been archived, models were re-run using *glmer.nb* in *lme4* package (version 1.1.27.1, Bates et al. 2015), which provides nearly identical parameter estimates and AICc values. The model fits were checked using the *DHARMa* package (version 0.4.6, Hartig 2022). Marginal means for genotype effects and their confidence intervals were calculated using the *emmeans* package (version, 1.8.5, Lenth 2023), whereby the values of continuous covariates were fixed at the mean value, and the value of the non-focal locus’ genotype was fixed at 0.5 (i.e., the EL genotype). We performed separate analyses for low- and high-nutrient streams to avoid overly complex model structures, since those two environments have been shown to have different associations with physiological indices (Auer et al. 2018).

### Partitioning additive and parental genetic effects

Given that the genotype effects for both *vgll3* and *six6* were significant in high nutrient streams (see Results below), we then employed individual level analyses to decompose the genetic variation to its additive (direct genetic effects) and parental genetic effects (indirect genetic effects), and further, to assess the contributions of each parent’s effects (i.e., maternal vs paternal, or both). Teasing out additive vs. parental genetic effects requires individual level survival data from the offspring, but we obviously did not have the genotype data for the missing (dead or emigrated) progeny. Therefore, the genotype of the missing progeny was inferred in accordance with Mendelian segregation of the parental genotypes, and after accounting for the known genotypes of the recaptured (surviving) individuals. As an example, suppose that for a family with parental *vgll3*^EE^ and *vgll3*^EL^ genotypes, nine offspring out of initial 100 implanted eggs survived in a particular stream, with seven of them being *vgll3*^EE^ and two of them being *vgll3*^EL^ genotype. In this case, and in accordance to Mendelian segregation laws, we can estimate that N=43 and N=48 offspring are missing (dead or emigrated) for the *vgll3*^EE^ and *vgll3*^EL^ genotype groups, respectively. A small subset of recaptured individuals with no genotype data (3.7% and 2.5% of the recaptured individuals for *six6* and *vgll3*, respectively) were randomly imputed a genotype based on Mendelian segregation of parental genotypes. (As an example, in a family with parental *vgll3*^EE^ and *vgll3*^EL^ genotypes, offspring with missing genotypes would be randomly assigned *vgll3*^EE^ or *vgll3*^EL^ genotypes with an equal (50%) probability). The dataset contained 29000 data points (100 offspring per 29 family and 10 streams), of which N=1246 consisted of recaptured individuals. The effect of sampling variation, due to both imputing missing genotypes of the recaptured individuals and missing individuals, was accounted for and quantified by resampling the missing genotypes 1000 times, which then re-analysed as described below.

We then built and compared the fit of binomial models with different parametrization of parental effects. All these models included family-wise SMR as a covariate (since this was significant in the high nutrient streams in the family-wise model – see Results), and the additive effects of the genotypes, but they differed in the parametrization of the parental effects, whereby each model contained a different combination of parental effects for *six6* and *vgll3* genotypes. As such, the general model structure could be written as follows:

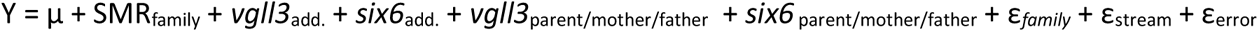

whereby, Y is a Boolean response variable for local survival (with missing and recaptured individuals scored as 0 and 1, respectively), SMRfamily is the family-wise mean SMR value, *vgll3*add. and *six6*add. are the additive genetic effects, and *vgll3*parent/mother/father and *six6* parent/mother/father are different combinations of parental effects associated with *six6* and *vgll3* loci. In total nine different parental genotype combinations were tested in nine models, whereby the parental covariates included in the models are as follows: (1) *vgll3*parent + *six6*parent, (2) *vgll3*parent + *six6*mother, (3) *vgll3*parent + *six6*father, (4) *vgll3*mother + *six6*parent, (5) *vgll3*mother + *six6*mother, (6) *vgll3*mother + *six6*father, (7) *vgll3*father + *six6*parent, (8) *vgll3*father + *six6*mother, (9) *vgll3*father + *six6*father. Family and stream were included in the models as random terms. The parental genotype value is the average genotype value of mother and father genotype, e.g., *vgll3*parent = (*vgll3*mother + *vgll3*father) / 2, and *six6*parent = (*six6*mother + *six6*father) / 2, which quantifies an overall combined parental genotype effect.

Genotypes were modelled additively (i.e., EE, EL, and LL alleles were coded as 0, 0.5, and 1, respectively) with no dominance effects for either genotype. The *vgll3* locus is known to have a sex-dependent dominance effect on sea-age at maturity, hence testing dominance in this locus would have provided further insights on the genetic architecture. However, the lack of *vgll3^LL^* genotypes in the parental fish and suited cross designs, we had one family with *vgll3^LL^*offspring (Supplementary Table 1), making predicting dominance effect unreliable. The fit of the models containing different combinations of parental genotypes were then compared according to the AICc criteria, in which models within ΔAICc < 2 to the best model were considered equally parsimonious.

To account for uncertainty associated with sampling variation of genotype distributions, we also performed random sampling of genotypes, in which genotypes of missing (dead or emigrated) individuals were randomly drawn from expected genotype probabilities, in accordance with Mendelian segregation of the parental genotypes. In this case, we simulated the genotypes of 100 offspring per family/stream (i.e., the initial number of individuals released per family to each stream) and assigned genotypes for missing individuals after excluding the genotypes of recaptured (surviving) individuals from the simulated set. Similar to above, a small subset of recaptured (survived) individuals with no genotype data (3.7% and 2.5% of the recaptured individuals for *six6* and *vgll3*, respectively) were also randomly assigned a genotype based on Mendelian segregation of parental genotypes in each replicate, to prevent any sampling biases. The random data sampling described here was repeated 1000 times.

## Results

The observed local survival per family was dependent on genotype in high but not low nutrient streams (see also the observed family-wise data in Supplementary Figure 1). In the low nutrient streams all parsimonious models (models within ΔAICc = 2 of the best model) contained egg mass and MMR as covariates, a result that is consistent with Auer et al. (2018). While the *vgll3* effect was present in the best model, it was insignificant, and the parameter was not present in all parsimonious models (Figure 1, Supplementary Table 2). In the high nutrient streams, all parsimonious models contained *six6* and SMR as covariates. In the best model, a higher SMR (*p* = 0.018) and *six6*^E^ allele (*p =* 0.025) were significantly and positively associated with local survival, while there was a marginal negative association with the *vgll3*^E^ allele (*p =* 0.047). Allelic substitution from *vgll3*^E^ to *vgll3*^L^ increased the odds of survival 1.67 times (SE = 0.432, *p* = 0.047), whereby the marginal survival probability associated with *vgll3*^EE^ was 3.16% (95% CI 2.46 -4.05), compared to 5.28% (95% CI 3.47 – 8.02) for *vgll3*^EL^ (Figure 1). Allelic substitution effect from *six6*^E^ to *six6*^L^ was in opposite direction, decreasing the odds of survival 1.49 times (SE = 0.118, *p* = 0.025), whereby the marginal survival probability associated with *six6*^EE^ was 7.85% (95% CI 4.94 -12.46), compared to 5.28% (95% CI 3.47 – 8.02) for *six6*^EL^ (Figure 1). Models were substantially more parsimonious when both genotype and physiological indices (SMR, MMR and egg mass) were included as covariates compared to the models that included only genotypes. However, family-wise SMR, MMR and egg mass were not significantly associated with either genomic region (Supplementary Table 3), suggesting an independent effect of genotypes and physiological traits on local survival rates. Similarly, an epistasis between *vgll3* and *six6* loci was not supported (Supplementary Table 3).

**Figure 1:**
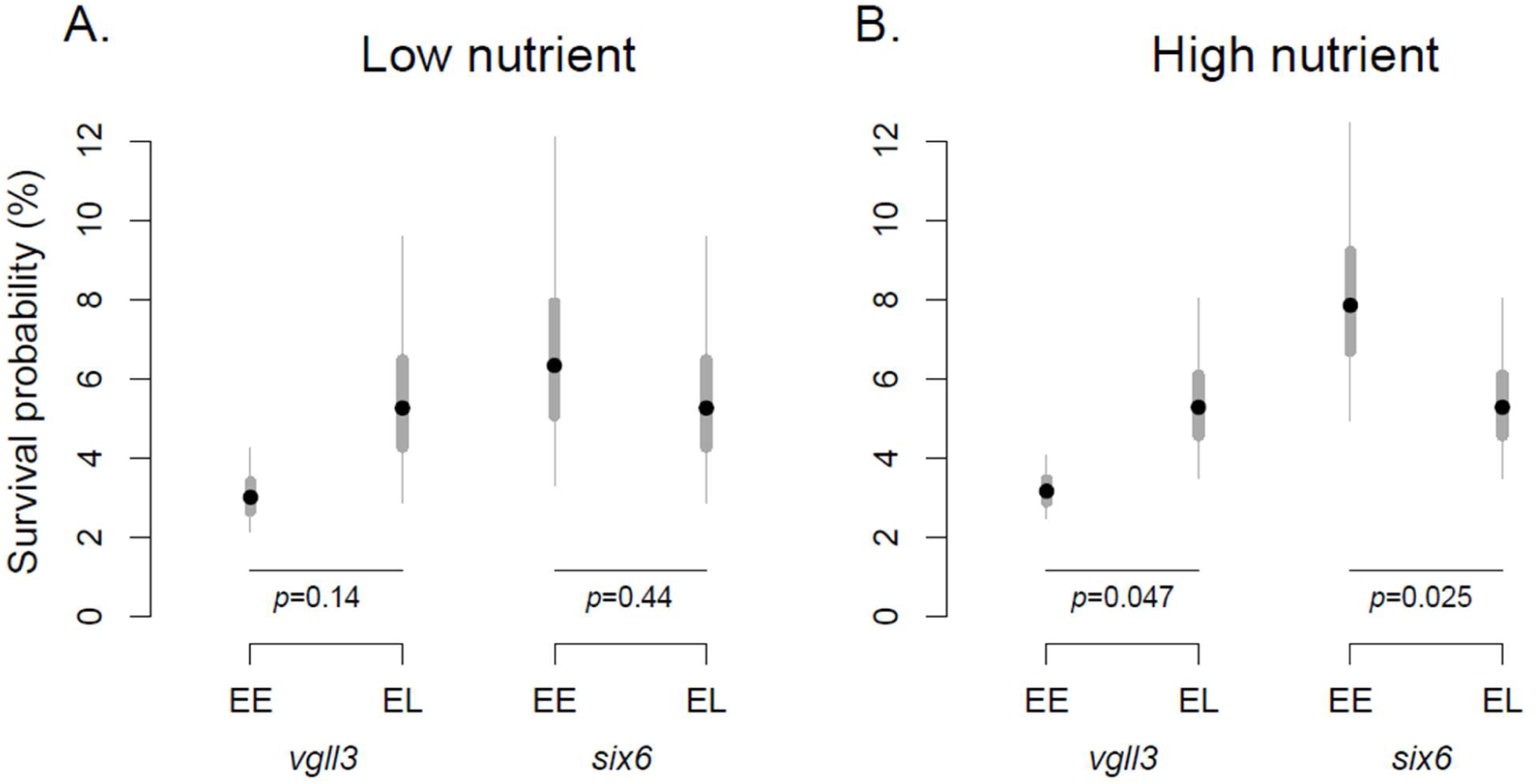
Marginal means for probability of survival of offspring in relation to family-wise *vgll3* and *six6* genotypes for (a) low, and (b) high nutrient streams. Data are plotted for the homozygote early (EE) and heterozygote (EL) genotypes (no homozygote-LL families were present for either locus). The estimated values are from the most parsimonious models including both *vgll3* and *six6* effects, which is the best fitting model for the high nutrient streams and the fourth best model for the low nutrient streams (see Materials and Methods and Supplementary Table 2 for details). Thin and thick grey bars indicate 50% and 95% confidence intervals, respectively.

For the high nutrient streams, we then used an individual-based, binomial model to partition the significant family-wise genotype effects into additive effects and different combinations of parental genetic effects. A model with only additive genetic terms had substantially lower fit compared to models that additionally parametrized parental genotypes as covariates (Figure 2). Among different combinations of parental genetic effects, the best model included parental *vgll3* and maternal *six6* genetic effects (Figure 2). This model was superior to the second-best model with an AICc differential 2.08 (SE=0.25, Figure 2). An allelic substitution from *vgll3*^E^ to *vgll3*^L^ in parental genotype increased the odds of survival by 1.98 times (SE = 0.60, *p* = 0.023), whereby the marginal survival probability associated with parental genotype *vgll3*^EE^ was 2.44% (95% CI 1.71 -3.46), compared to 4.72% (95% CI 3.09 – 7.15) for *vgll3*^EL^ (Figure 3, Table 1). Likewise, an allelic substitution from *six6*^E^ to *six6*^L^ associated with maternal genotype decreased the odds of survival by 1.58 times (SE = 0.10, *p* = 0.003), whereby the marginal survival probability associated with *six6*^EE^ maternal genotype was 7.25% (95% CI 4.69 - 11.07), compared to 4.72 % (95% CI 3.09 – 7.15) for *six6*^EL^ (Figure 3). The additive (direct) *E* to *L* allelic substitutions was insignificant both for *vgll3* (*p =* 0.51) and *six6* loci (*p =* 0.36, Figure 3, Table 1). The individual-based analyses were robust to sampling variation associated with imputing the missing genotypes of missing individuals, or recaptured individuals that were not successfully genotyped, since similar effect sizes were obtained across 1000 simulated datasets (Supplementary Figure 2). The results were also robust to some families being invariant in *vgll3* and *six6* genotypes, which may result in lower power for quantifying the additive genetic variance compared to the parental genetic effects. As such, when the analyses included only families which harboured genetic variation in these loci (21 and 14 families for *vgll3* and *six6*, respectively, see Supplementary Table 3), the effect sizes and associated *p-values* were similar to those obtained from the full dataset, and parental genetic effect sizes were substantially higher than the additive genetic effects (Supplementary Table 4).

**Figure 2:**
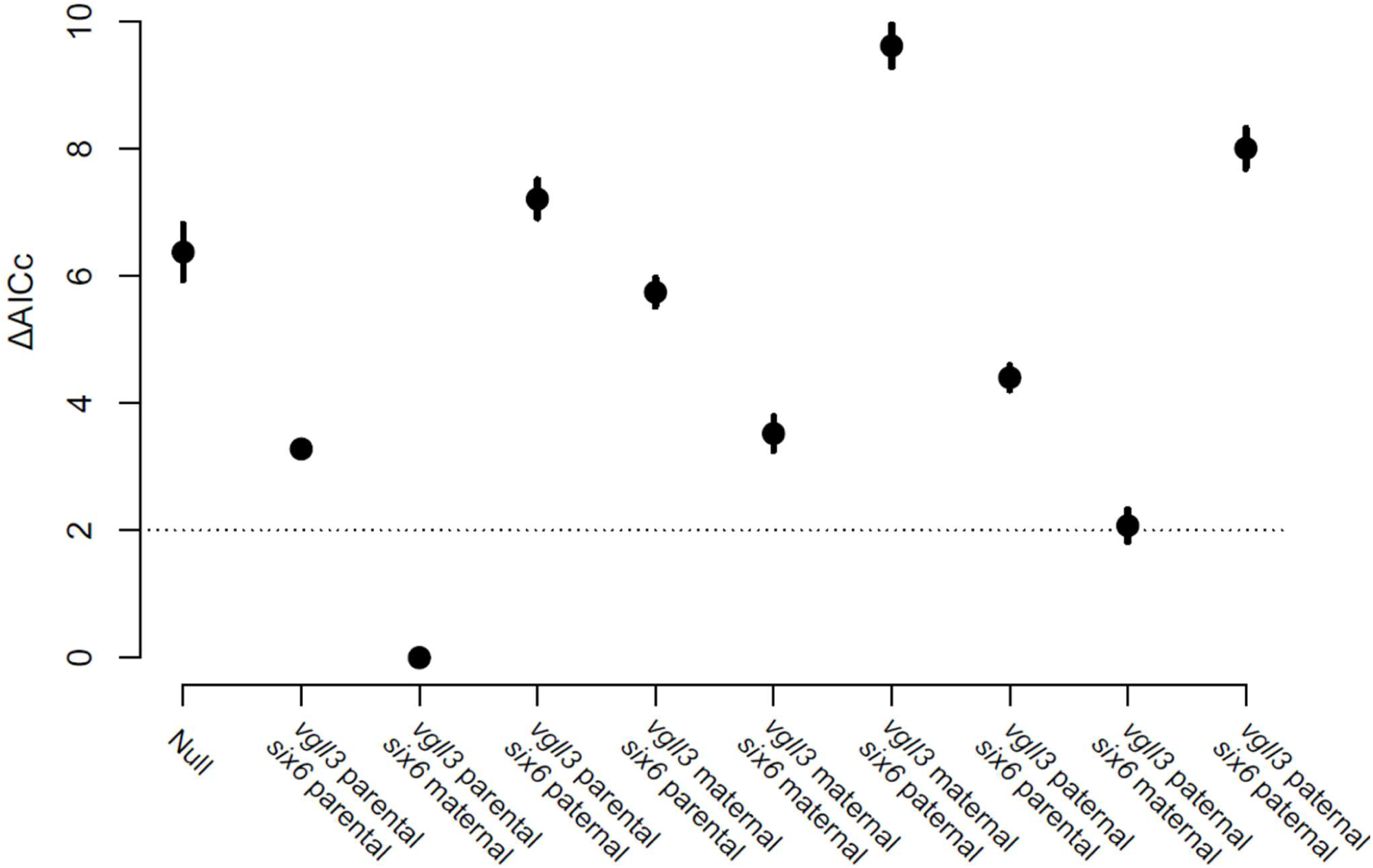
The likelihood differences among individual-based models with different parameterization of the parental genetic effects explaining survival in the high nutrient environment. ΔAICc = 0 indicates the best fitted model. All models contained SMR and additive *vgll3* and *six6* genetic effects as covariates. Y-axis indicates the ΔAICc to the best model, and the dashed line indicates ΔAIC = 2. Error bars represents one standard error of ΔAICc across 1000 models using 1000 data that simulated the genotypes of dead/emigrated individuals (see Materials and Methods for details).

**Figure 3:**
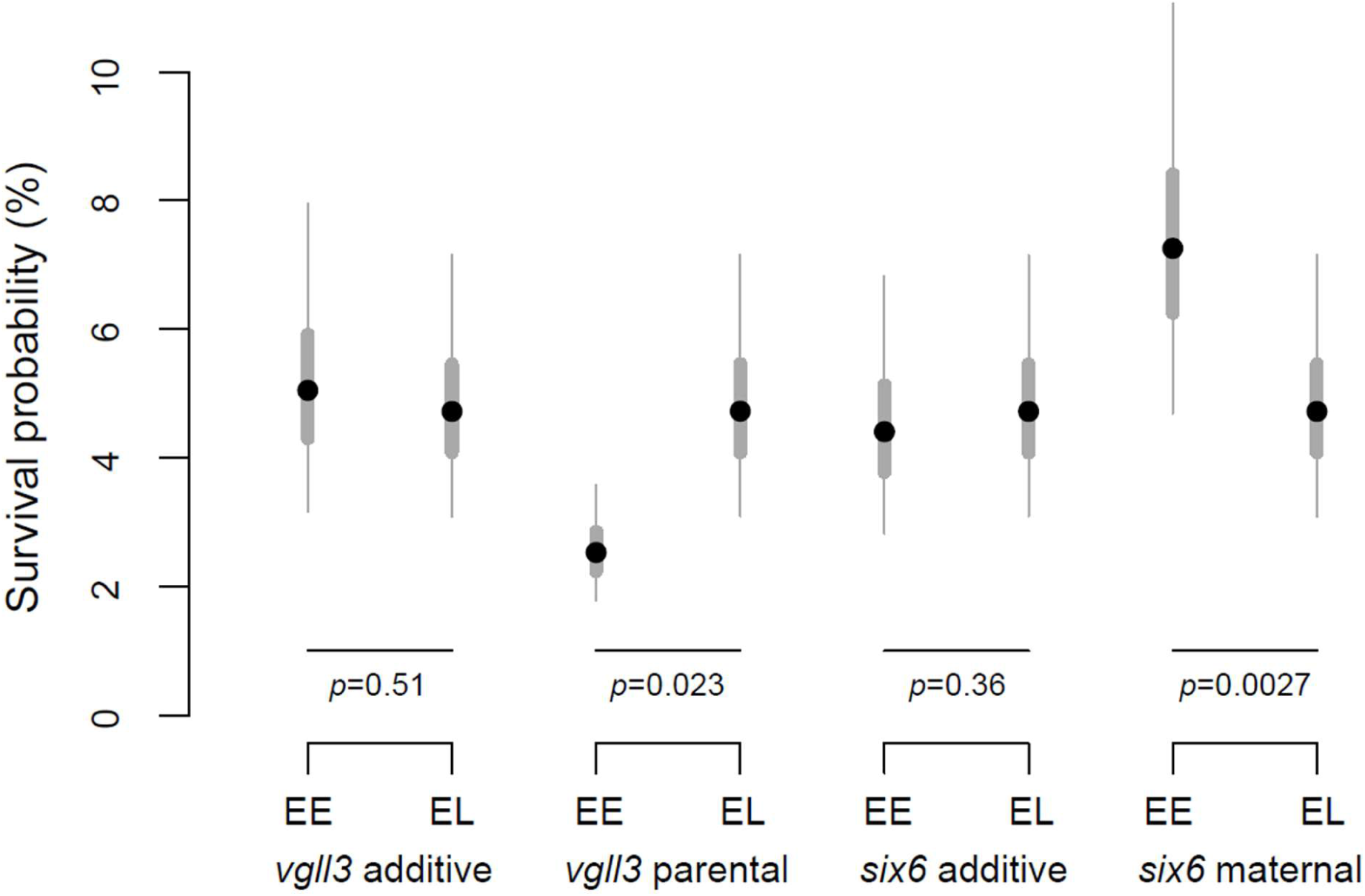
Genetic effects on marginal offspring survival probability for the best-fitted parental genetic structure model in the high nutrient streams. Thin and thick grey bars indicate 50% and 95% confidence intervals, respectively. Data are plotted for the homozygote early (EE) and heterozygote (EL) genotypes (no homozygous-LL families were present for either locus).

**Table 1:** Estimated coefficients (fixed effects) and variance (random effects) for individual based binomial model with local survival as the response variable. The coefficients are from the best-fitted model across different parametrization of the parental genetic effects in the high nutrient environment.

| **Fixed effects** | Estimate | Std. Error | z value | p value |
| --- | --- | --- | --- | --- |
| (Intercept) | -5.959 | 1.156 | -5.154 | <0.0001 |
| Family SMR | 16.142 | 6.945 | 2.324 | 0.0201 |
| *vgll3* additive | -0.141 | 0.213 | -0.662 | 0.5080 |
| *six6* additive | 0.145 | 0.159 | 0.911 | 0.3625 |
| *vgll3* parental | 1.368 | 0.602 | 2.274 | 0.0230 |
| *six6* maternal | -0.914 | 0.305 | -3.002 | 0.0027 |
| **Random effects** | Variance |  |  |  |
| Family origin | 0.1543 |  |  |  |
| Stream | 0.0047 |  |  |  |

In salmonids, the maternal genotype might be more important to offspring survival than the paternal genotype, since mothers also determine the egg size and composition that influences offspring viability (Fleming 1996). To test the if maternal body size drives the association between local survival and the maternal genetic effects (in *six6* locus), maternal body size traits (i.e., length, weight, and condition factor) were added separately to the most parsimonious models explaining the maternal genotype effects on offspring local survival in the high nutrient streams. None of the maternal size traits had a significant effect on survival and did not improve the model fit, with or without the *six6* maternal effects present in the models (Supplementary Table 5). This suggests that maternal size is not driving the association between offspring survival and maternal *six6* genotypes, although it should be noted that variation in maternal size within the sample was constrained as a result of only using females that had spent a single winter at sea. On the other hand, a maternal *six6* genotype had strong associations with her size and condition (Supplementary Table 6), with the late genotype being associated with larger size and higher body condition. This indicates that a genetic correlation exists between parental size and early offspring survival via *six6*, but the association between *six6* and offspring survival could be mediated by a different process than female body size.

## Discussion

Understanding the genetic architecture of adaptation is crucial for predicting how populations respond to changing environmental conditions at contemporary time scales. Major effect loci may alter phenotypes at early life stages (i.e., during development), leading to phenotypic and fitness changes operated via multiple pathways (Colosimo et al. 2005). This study has demonstrated that the loci that are associated with maturation timing in Atlantic salmon can also affect early life survival, but in a manner that interacts with nutrient levels and the availability of food in the streams that they inhabit. Intriguingly, this effect did not operate via additive genetic effects associated with offspring own genotype, but via a maternal effect in *six6* loci, and via parental effects in *vgll3*. The results suggest that the offspring of parents carrying the *vgll3* late-maturation genotype and of mothers carrying the *six6* early-maturation genotype will benefit from higher survival, regardless of the inherited alleles. Such indirect genetic effects add further complexity to evolutionary dynamics compared to simple Mendelian inheritance (Bonduriansky and Day 2009; Wolf et al. 1998), but have been little studied in wild populations.

Salmon are well known for exhibiting plasticity in life history traits, particularly in response to environmental conditions that influence growth rates (Mobley et al. 2021; Sloat et al. 2014). Age at maturity is an important life history trait that can influence fitness and that has a genetic as well as an environmental component, likely maintained in the population via balancing selection (Barson et al. 2015). Our results suggest that the viability selection acting on *vgll3* and *six6* loci in early juvenile stages of salmon will likely alter the age-structure, hence the dynamics of the evolutionary response in the population. However, the response has a plastic component and operates in environments with higher nutrient levels. As such, the rate of evolutionary response in age at maturity may be attributed, in part, to ecological processes operating very early in life. However, the effects of the freshwater environment on variation in age at maturity are poorly understood, limiting the scope for *a priori* predictions of the effect of these life-history loci on early survival (see also the Introduction). Similarly, while our results indicate that a genetic correlation exists between early- and late-life fitness components, it does not reveal the possible physiological basis or how the observed correlation between fitness components contributes to the complexity of adaptation (Orr 2000). For example, it is unclear if the direction of selection associated with early- and late-life fitness components is aligned (i.e., synergistic pleiotropy) or inflicts a cost to the life-history adaptations (i.e., constraining selection via trade-offs, Orr 2000; Rennison and Peichel 2022; Thorholludottir et al. 2023). A recent study showed that a rapid evolutionary response towards earlier maturation (i.e., younger age structure) in the *vgll3* locus was partly driven by the abundance of prey species once the salmon reach the ocean (Czorlich et al. 2022). However, our results in the high nutrient streams suggest that selection in earlier life stages favours the opposite *vgll3* allele (i.e., late). One might then predict a larger evolutionary cost to age-at-maturity evolution at the *vgll3* loci, i.e., via a slower evolutionary response to a changing environment (and a larger genetic load to the population, Orr 2000). However, we found that the genetic effects on survival was dependent on nutrient levels, and therefore the potential for correlated selection with age at maturity is context dependent, suggesting that the cost of any correlation between fitness components might be population/condition dependent. Finally, the survival effects associated with *six6* and *vgll3* loci are in opposite directions to each other with respect to their effect on maturation timing: while the *vgll3*^L^ allele (the allele linked to later age at maturity) appears to be associated with higher offspring survival, it is the early maturation allele (*six6*^E^) that positively influences offspring survival in the *six6* region, which necessitates that variation in both loci is accounted for when formulating predictions. Put together, our results highlight the complexity related to predicting the evolutionary dynamics of maturation age in Atlantic salmon, and potentially in other taxa where indirect genetic effects have been found (e.g., Bonduriansky and Day 2009; Tomar and Teperino 2020, Wolf et al. 1998). The observed genetic correlation, however, suggests that a physiological, mechanistic link exists between genetic polymorphisms in the life-history loci and early survival.

The indirect genetic effect on viability selection via parental genotypes might have theoretical implications for evolutionary processes (e.g., Bonduriansky and Day 2009; Rausher 1992; Räsänen and Kruuk 2007). In the classical sense, parental effects are a component of phenotypic plasticity, in which the parental environment influence the offspring phenotype, through developmental reprogramming (Kappeler and Meaney 2010). As such, indirect genetic effects via parental genetic variation help to maintain genetic diversity i.e., by reducing the narrow sense heritability by increasing environmental variance, and provide a means for transmission of acquired traits between generations (Bonduriansky and Day 2009; Wolf et al. 1998). However, in our context, the indirect genetic effects of parents are also correlated with the offspring’s future life-history and thereby with the fitness of the offspring via the direct effect of genotypes on age at maturity (Wolf et al. 1998). Therefore, selection on offspring survival would alter the phenotypic variation of life-history traits, resulting in deviating from the optimal trait variation maintained under balancing selection (Turelli and Barton 2004), though this prediction is based on a relatively limited range of genotypes (since the *vgll3*^LL^ genotypes was completely absent in the dataset) and phenotypes (since all parents were 1SW fish). Theoretical predictions accounting for such a complexity have not been explored explicitly (e.g., Oomen et al. 2020).

Our results highlight that the effect of viability selection on these genomic regions is dependent on the environment, as it appears to only operate under conditions of high nutrient availability. Both loci are linked to a number of different physiological and behavioural traits that mediate energy homeostasis (Mobley et al. 2021). However, we did not find a direct link between these genomic regions and SMR or MMR (measured at the family level), thereby suggesting that metabolic rates are not underlying mechanisms for mediating the effects of *vgll3*/*six6* loci on early survival. Earlier studies have shown a broad range of effects on physiological and behavioural traits linked to genetic variation in *vgll3* locus, such as MMR (Prokkola et al. 2022), lipid profiles (House et al. 2021), and aggressive behaviour (Bangura et al. 2022), and that the *six6* genomic region can be broadly expressed during embryonic developmental (Moustakas-Verho et al. 2020). The effect of these on fitness is likely to be context-dependent, supporting our conclusions, although the exact mechanisms are yet to be elucidated. In our study, the genotype dependent selection operating under a more favourable environment (high nutrient) may seem counterintuitive. However, the higher relative fitness (of different genotypes) in a less stressful environment is not only theoretically possible, but an earlier meta-analysis indicated no overall differences in relative fitness under more stressful conditions (Arbuthnott and Whitlock, 2018). Furthermore, there were no differences in absolute fitness between low- and high-nutrient conditions, despite substantial differences in juvenile growth rates and different metabolic processes shaping the fitness landscape (Auer et al. 2018), and different impacts of habitat features on the rate of senescence in the fish (McLennan et al. 2021). These suggests a complex landscape is likely underlying the fitness dynamics across environment.

The higher local survival of juvenile fish with the early maturation genotype at the *six6* locus was explained solely by the maternal genotype. Indirect genetic effects associated with the maternal environment (i.e., maternal effects) are commonly observed in salmonids and in other organisms (e.g., Aykanat et al. 2012; Heath et al. 1999; Mousseau and Fox 1998). However, because salmonids lack parental provisioning, these effects are mostly limited to egg size and/or composition, which in turn can be influenced by the size of the mother (Munoz et al. 2014). A significant trait-genotype association (i.e., between *six6*mother and offspring survival / mother size) despite that the mother’s size was constrained (due to using only females that spent a single winter at sea), suggests a robust inference can be drawn to test for a potential effect of an intermediate phenotype (egg mass) mediating the relation. Our results showed that in the high nutrient streams, family-level egg mass was neither correlated with survival (Supplementary Table 3), nor with variation in the *six6*mother (Supplementary Table 2), thereby ruling out a genetic effect of egg mass on survival via *six6*mother (within the constrained size range of mothers). Similarly, we found that maternal size did not explain the correlation between *six6*mother and the survival of her offspring, despite the observed correlation between maternal size and *six6*mother (Supplementary Table 6).

A plausible mechanism explaining our results is that that variation in *six6* alters early life survival via changes in egg composition. In oviparous species, the composition of eggs, such as hormone levels, albumin and fat content, may mediate the phenotype and behaviour of the offspring, and subsequently their fitness (Thorn and Morbey 2019; Widowski et al. 2022; Williams 1994). In Chinook salmon, carotenoid composition in the eggs has been shown to be associated with early life survival (Tyndale et al. 2008). Therefore, despite the apparent absence of a link between the *six6* region and family-level egg mass, we cannot rule out potential effects on offspring survival via changes in egg composition. Future studies could test such a hypothesis in a common garden cross design, by quantifying potential markers associated with egg composition, and their effect on the survival of their full sibs. Alternatively, but less parsimoniously, epigenetic mechanisms may be associated with a mothers’ *six6* genotype, whereby the maternal *six6* locus controls the epigenetic signature of the offspring genome and thus influences early life developmental pathways (see, Lawson et al. 2013; Venney et al. 2021; Wolf and Wade 2009). *Six6* and other candidate genes located in the same genomic region (i.e., *six1* and *six4*) are known transcription factors, and *six6* expression has been documented in Atlantic salmon in early development (Moustakas-Verho et al. 2020). It is therefore plausible that a mother’s *six6* genotype may play a role in the regulation of her offspring’s genome.

The effect of a parent’s *vgll3* genotype on offspring early life survival was not solely limited to the maternal contribution. In salmonids, and generally in fishes, the lack of parental provisioning rules out social environment such as behavioural variation as a potential basis of indirect genetic effects (e.g., Bailey et al. 2018, Bonduriansky and Day 2009). A plausible mechanism that is consistent with the observed results is that the genomes inherited from parents are already modified via epigenetic mechanisms, such as via DNA methylation (Bonduriansky and Day 2009; Kappeler and Meaney 2010; Lawson et al. 2013). Epigenetic regulation is known to alter phenotypic variation in natural populations, but the underlying genetic mechanisms and evolutionary predictions are still unclear (Husby 2022). In salmonids, epigenetic effects via DNA methylation have been documented, for example, between hatchery and wild origin fish associated with both parents of origin, with implications for evolutionary genetics and conservation efforts (Koch et al. 2022; Le Luyer et al. 2017; Smith and Kornfield 2002). Our results are consistent with a hypothesis whereby variation in epigenetic signals (associated with genetic variation in the *vgll3* locus) could result in differences in gene expression during early development, subsequently altering offspring survival in the natural settings. Such a hypothesis may be tested by designing cross-breeding experiments, where offspring epigenetic signals, such as genome wide methylation patterns, could be examined among crosses from parents with different *vgll3* genotypes, and in relation to offspring phenotype and survival (i.e., Bossdorf et al. 2008). Alternatively, histone modification is another plausible mechanism that might generate parental genotype-specific modification of chromatin structure, and subsequent survival differences between families (Koch et al. 2022).

Even though we have shown that the same genomic regions can account for variance in different fitness components, it is unclear whether covariation between fitness components is causally linked to a single genetic mutation (true pleiotropy), or takes place as a result of physically linked loci acting on different fitness components. While molecular evidence suggests that both *vgll3* and *six6* regulate multiple functions across multiple life stages (Mobley et al. 2021), implying pleiotropy, genetic linkage within the genomic regions cannot be ruled out. There is accumulating evidence to suggest that co-adapted gene complexes are common in wild species (Li et al. 2022; Matschiner et al. 2022; Thompson and Jiggins 2014). Moreover, theoretical models have predicted the evolution of genetic architecture with fewer, larger, and more tightly linked divergent alleles in species that exhibits marked local adaptation between migratory populations (Yeaman and Whitlock 2011). In keeping with this model, Atlantic salmon exhibit local adaptation across closely situated populations with both *six6* and *vgll3* genomic regions harbouring substantial signals of divergence, even between adjacent populations (e.g., Barson et al. 2015; Pritchard et al. 2018). Therefore, it is plausible that the genetic correlations of fitness components via the *six6* and *vgll3* genomic regions might be explained by independent loci with tight linkage within these genomic regions. Such genetic architecture may be instrumental in maintaining the genetic variation and local adaptation of populations amidst high gene flow, although explicit mathematical models and empirical tests of this hypothesis is still lacking.

Our study re-purposed a dataset from Auer et al. (2018) in order to test the effect of life history genomic regions on early life survival. While such approaches provide a substantial savings in resources, they come with unavoidable drawbacks. In particular, we could not explicitly test for dominance effects in the *vgll3* locus, which exhibits a (sex-dependent) dominance architecture associated with the age-at-maturity genotype (Barson et al. 2015). This is because the genotypes in the parental fish were either *vgll3^EE^* or *vgll3*^EL^, and only one family (out of 29) had a parental genotype combination that would generate *vgll3*^LL^ offspring (i.e., both parents heterozygous for *vgll3*), making inferences about dominance confounded with the additive effect, and thus unreliable. For *six6*, we did not have a priori belief that dominance should explain the trait value, and it was not considered in the main framework. However, modelling dominance in the *six6* locus did not improve the model fit (AICcdominance – AICcadditive = 2.00), and the *six6* effect was still insignificant, as it is when modelled additively.

In this study, we provide a unique example from a natural setting of how indirect genetic effects from large-effect loci influencing adult age at maturity and adaptive divergence shape the early-life survival of offspring. Indirect genetic effects are common across organisms and can alter phenotypic variation and evolutionary trajectories, but most empirical evidence is based on transmission of behavioural traits in higher organisms, or has explored typical maternal effects (Wolf et al. 1998; Bonduriansky and Day 2009). The complex genetic architecture of large effect loci that comprise indirect genetic effects between parents and offspring revealed by this study highlights the challenges associated with predicting evolutionary dynamics in a changing world. Little is known of the fitness landscapes of large effect loci across life stages in the wild, and the generality of our results across different Atlantic salmon populations and taxa needs further work. Our study also calls for a better understanding of the mechanisms, ecological role and evolutionary consequences of indirect genetic effects.

## Data accessibility

The compiled datasets and R-codes to reproduce the results are available from the dryad repository https://doi.org/10.5061/dryad.866t1g1xk.

## Acknowledgements

We thank the Cromarty Firth Fishery Trust (in particular the late Simon McKelvey), Sonya Auer and our field team for assistance in fieldwork, and Leena Laaksonen and the MES Lab in the University of Helsinki for help with the KASP assays. Funding came from the European Research Council (Advanced Grants 322784 and 834653 to NBM), and from the Research Council of Finland (353388 and 325964 to TA, 348965 to JMP).

