## Supplementary Materials for "Early survival in Atlantic salmon is associated with parental genotypes at loci linked to timing of maturation"

for

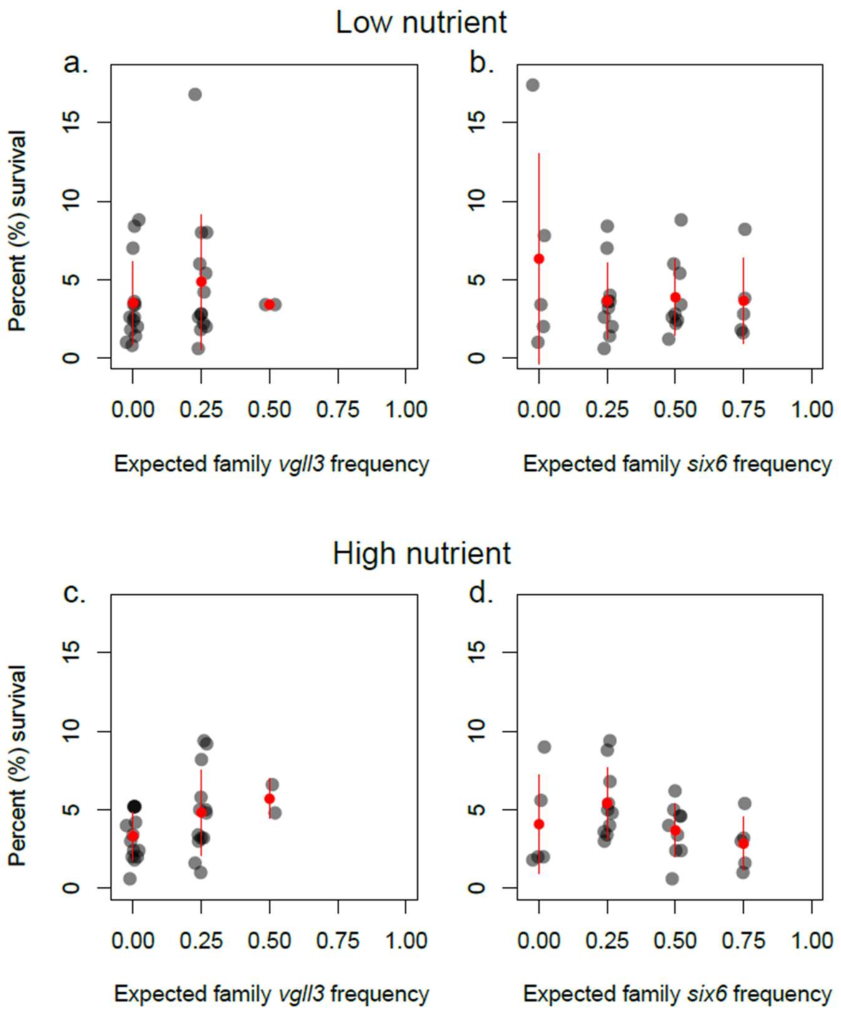


**Supplementary Figure 1:** Percent (%) local survival of each of 29 families of Atlantic salmon as a function of *vgll3* and *six6* genotypes, in low (a, b) and high (c, d) nutrient conditions, averaged across five streams per nutrient condition. Numbers on the x-axis indicate the expected average gene frequency of each full-sib family under Mendelian segregation, where 0 indicates both parents were homozygous for the early, and 1 indicates both parents are homozygous for the late allele (hence all progeny would also be homozygous). Data points are slightly jittered along the X-axis to improve visibility. Red points and lines indicate the group means and the standard deviation.

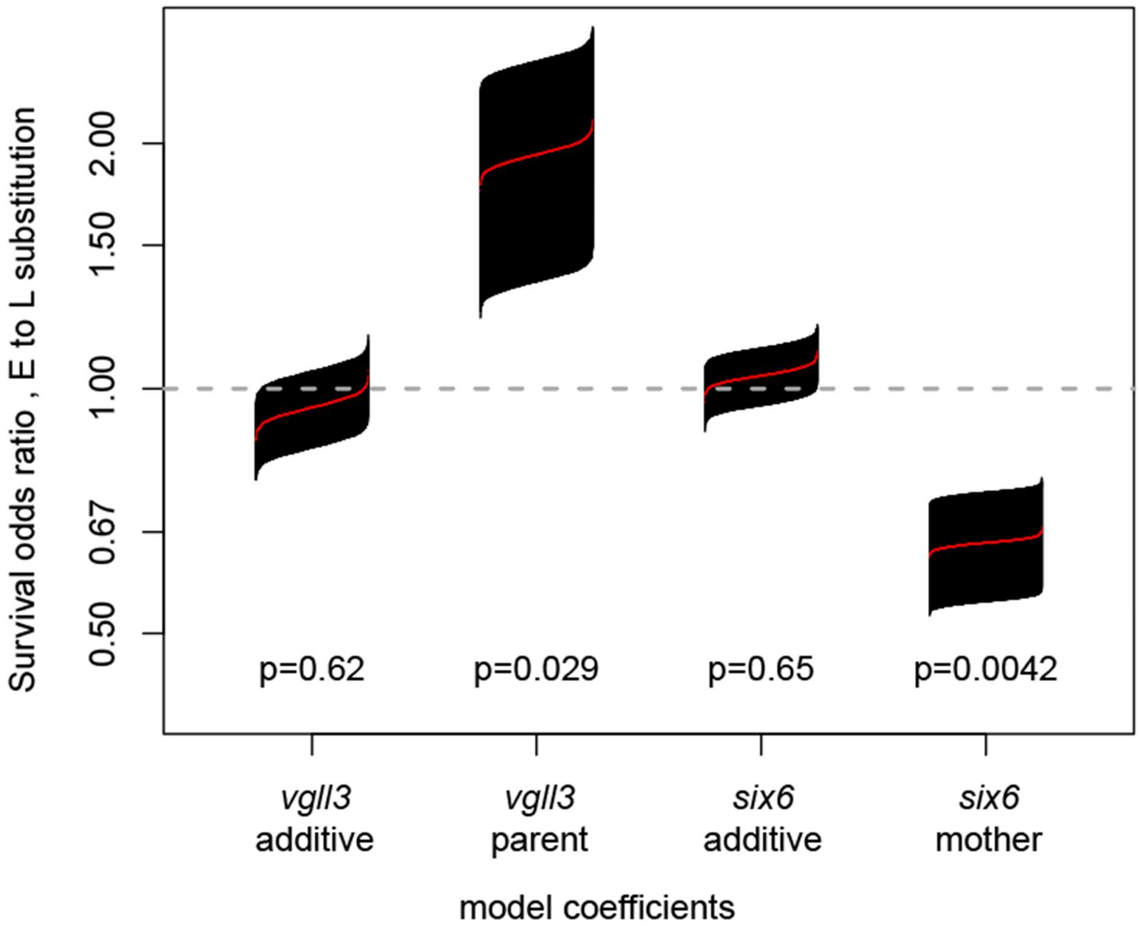


**Supplementary Figure 2:** Genotype dependent odds-ratios of survival from the individuals-based model with the most parsimonious parental genetic structure from 1000 models that used 1000 simulated genetic data for dead/emigrated individuals . Red line shows sorted mean estimates from 1000 models, while black area indicates the standard errors. Dashed line indicates no effect value (OR =1). Y-axis is plotted on log scale to better visualise effect-size in increasing (above 1) vs decreasing (below 1) order. P-values are averaged across 1000 models.

**Supplementary Table 1**: The genetic structure for *vgll3* and *six6* loci across 29 families used in the study. EE and LL homozygote genotypes corresponds to 0 and 1, respectively, and EL heterozygote genotype corresponds to 0.5.

| Family ID | Maternal genotypes | | Paternal genotypes | | Expected family-wise *genotype* frequency | | Expected number of genotypes | |
|---|---|---|---|---|---|---|---|---|
| | *vgll3* | *six6* | *vgll3* | *six6* | *vgll3* | *six6* | *vgll3* | *six6* |
| 1 | 0 | 0.5 | 0 | 1 | 0 | 0.75 | 1 | 2 |
| 2 | 0 | 0 | 0 | 0.5 | 0 | 0.25 | 1 | 2 |
| 3 | 0 | 0 | 0 | 0 | 0 | 0 | 1 | 1 |
| 4 | 0 | 0 | 0.5 | 0.5 | 0.25 | 0.25 | 2 | 2 |
| 5 | 0 | 0 | 0 | 0.5 | 0 | 0.25 | 1 | 2 |
| 6 | 0 | 1 | 0 | 0 | 0 | 0.5 | 1 | 1 |
| 7 | 0 | 0 | 0 | 0 | 0 | 0 | 1 | 1 |
| 8 | 0 | 1 | 0.5 | 0.5 | 0.25 | 0.75 | 2 | 2 |
| 9 | 0.5 | 0.5 | 0.5 | 0.5 | 0.5 | 0.5 | 3 | 3 |
| 10 | 0.5 | 0.5 | 0 | 0.5 | 0.25 | 0.5 | 2 | 3 |
| 11 | 0 | 0.5 | 0 | 1 | 0 | 0.75 | 1 | 2 |
| 12 | 0 | 0 | 0.5 | 0.5 | 0.25 | 0.25 | 2 | 2 |
| 13 | 0 | 0 | 0 | 0 | 0 | 0 | 1 | 1 |
| 14 | 0.5 | 0.5 | 0 | 0 | 0.25 | 0.25 | 2 | 2 |
| 15 | 0 | 0 | 0.5 | 0.5 | 0.25 | 0.25 | 2 | 2 |
| 16 | 0.5 | 0 | 0 | 0 | 0.25 | 0 | 2 | 1 |
| 17 | 0 | 0 | 0 | 0.5 | 0 | 0.25 | 1 | 2 |
| 18 | 0 | 0 | 0.5 | 0.5 | 0.25 | 0.25 | 2 | 2 |
| 19 | 1 | 0 | 0 | 0.5 | 0.5 | 0.25 | 1 | 2 |
| 20 | 0 | 0 | 0 | 0.5 | 0 | 0.25 | 1 | 2 |
| 21 | 0 | 0.5 | 0 | 0.5 | 0 | 0.5 | 1 | 3 |
| 22 | 0 | 0 | 0 | 1 | 0 | 0.5 | 1 | 1 |
| 23 | 0 | 0.5 | 0.5 | 0.5 | 0.25 | 0.5 | 2 | 3 |
| 24 | 0 | 0 | 0.5 | 0 | 0.25 | 0 | 2 | 1 |
| 26 | 0.5 | 0.5 | 0 | 1 | 0.25 | 0.75 | 2 | 2 |
| 27 | 0 | 0 | 0 | 1 | 0 | 0.5 | 1 | 1 |
| 28 | 0 | 0.5 | 0 | 1 | 0 | 0.75 | 1 | 2 |
| 29 | 0.5 | 0.5 | 0 | 0.5 | 0.25 | 0.5 | 2 | 3 |
| 30 | 0 | 0.5 | 0.5 | 0.5 | 0.25 | 0.5 | 2 | 3 |

**Supplementary Table 2:** Generalized linear mixed models with negative binomial error structure predicting local survival in low and high nutrient treatments. Candidate models included other physiological indices (MMR, SMR, and egg mass) as fixed terms in addition to *six6* and *vgll3* genotypes and their interaction. Only the most parsimonious models (dAICc < 2 to the best model), and the following best model are included in the table. All models contained stream and family as random terms. A cell is left empty if the term is not included in the particular model.

| | **Intercept** | **egg mass** | **MMR** | **SMR** | ***six6*** | ***vgll3*** | ***six6*:*vgll3*** | **df** | **AICc** | **d(AICc)** |
|---|---|---|---|---|---|---|---|---|---|---|
| **Low Nutrient** | -9.595 | 0.019 | 18.260 | | | 1.167 | | 7 | 719.81 | 0.00 |
| | -9.101 | 0.018 | 17.890 | | | | | 6 | 720.07 | 0.26 |
| | -8.384 | 0.019 | 16.520 | | -0.442 | | | 7 | 721.45 | 1.64 |
| | -9.046 | 0.020 | 17.220 | | -0.372 | 1.119 | | 8 | 721.45 | 1.64 |
| | -9.351 | 0.020 | 18.420 | -2.378 | | 1.237 | | 8 | 721.99 | 2.19 |
| **High Nutrient** | -1.172 | | | 16.113 | -0.794 | 1.026 | | 7 | 739.72 | 0.00 |
| | -0.926 | -0.006 | | 17.960 | -0.727 | 0.987 | | 8 | 740.82 | 1.10 |
| | -1.451 | | | 18.687 | -0.834 | | | 6 | 741.01 | 1.29 |
| | -1.095 | | | 15.150 | -0.565 | 1.767 | -2.014 | 8 | 741.34 | 1.62 |
| | -1.146 | -0.007 | | 20.831 | -0.760 | | | 7 | 741.83 | 2.11 |

**Supplementary Table 3:** Linear models for family-wise metabolic traits in relation to each other, and family-wise *six6* and *vgll3* genotype (and their interaction) as measured in the common garden settings (Auer et al 2018). Only the most parsimonious models (dAICc < 2 to the best model), and the following best model are included in the table. A cell is empty if the term is not included in a model.

| Trait | intercept | egg mass | SMR | MMR | *six6* | *vgll3* | *six6*:*vgll3* | df | AICc | d(AICc) | weight |
|---|---|---|---|---|---|---|---|---|---|---|---|
| **SMR** | 0.1432 | 0.00027 | NA | | | | | 3.00 | -163.34 | 0.00 | 0.24 |
| | 0.1393 | 0.00027 | NA | | | 0.02380 | | 4.00 | -163.11 | 0.24 | 0.21 |
| | 0.1688 | | NA | | | | | 2.00 | -162.97 | 0.37 | 0.20 |
| | 0.1654 | | NA | | | 0.02341 | | 3.00 | -162.63 | 0.71 | 0.17 |
| | 0.1647 | | NA | | 0.01103 | | | 3.00 | -161.64 | 1.70 | 0.10 |
| | 0.1424 | 0.00024 | NA | | 0.00818 | | | 4.00 | -161.31 | 2.03 | 0.09 |
| **MMR** | 0.4881 | | | NA | | | | 2.00 | -137.02 | 0.00 | 0.56 |
| | 0.4935 | | | NA | -0.01459 | | | 3.00 | -135.35 | 1.67 | 0.24 |
| | 0.4905 | | | NA | | -0.016 | | 3.00 | -134.92 | 2.10 | 0.20 |
| **Egg mass** | 36.40 | NA | 351.10 | | | | | 3.00 | 245.15 | 0.00 | 0.39 |
| | 95.67 | NA | | | | | | 2.00 | 245.51 | 0.37 | 0.33 |
| | 91.33 | NA | | | 11.69 | | | 3.00 | 247.02 | 1.87 | 0.15 |
| | 38.28 | NA | 322.14 | | 8.14 | | | 4.00 | 247.35 | 2.20 | 0.13 |

**Supplementary table 4:** Estimated coefficients (fixed effects) for individual based binomial models with local survival as the response variable obtained from the reduced datasets, in which only families that are variable for *vgll3* (N=14) and *six6* (N=21) is implemented. The models are the best-fitted model across different parametrization of the parental genetic effects in the high nutrient environment.

| | **Fixed effects** | Estimate | Std. Error | *z* value | *p* value |
|---|---|---|---|---|---|
| Reduced dataset for *vgll3* (N=14 families) | (Intercept) | -1.293 | 2.707 | -0.478 | 0.6330 |
| | Family SMR | -11.157 | 15.121 | -0.738 | 0.4606 |
| | *vgll3* additive | -0.101 | 0.219 | -0.460 | 0.6457 |
| | *six6* additive | 0.160 | 0.202 | 0.790 | 0.4293 |
| | *vgll3* parental | 2.105 | 1.431 | 1.471 | 0.1413 |
| | *six6* maternal | -1.360 | 0.465 | -2.923 | 0.0035 |
| | **Random effects** | Variance | | | |
| | Family origin | 0.178748 | | | |
| | Stream | 0.003424 | | | |
| Reduced dataset for *six6* (N=21 families | (Intercept) | -3.554 | 1.364 | -2.606 | 0.0092 |
| | Family SMR | 2.785 | 7.829 | 0.356 | 0.7221 |
| | *vgll3* additive | -0.079 | 0.239 | -0.332 | 0.7397 |
| | *six6* additive | 0.221 | 0.165 | 1.334 | 0.1821 |
| | *vgll3* parental | 1.282 | 0.531 | 2.416 | 0.0157 |
| | *six6* maternal | -1.166 | 0.291 | -4.012 | 0.0001 |
| | **Random effects** | Variance | | | |
| | Family origin | 0.07218 | | | |
| | Stream | 0.01028 | | | |

**Supplementary table 5:** Comparing maternal *six6* effect and maternal size variables to early-life survival by model comparisons between best parental genetic effects model in the high nutrient streams (as in Table 1) to the models that additionally includes maternal (mother) length, weight, and condition, separately. All models contained stream and family as random terms.

| **Model** | **df** | **AICc** |
|---|---|---|
| SMR + *vgll3* additive + *six6* additive + *vgll3* parental + *six6* mother | 8 | 5022.994 |
| SMR + *vgll3* additive + *six6* additive + *vgll3* parental + *six6* mother + length.mother | 9 | 5024.959 |
| SMR + *vgll3* additive + *six6* additive + *vgll3* parental + *six6* mother + weight.mother | 9 | 5024.71 |
| SMR + *vgll3* additive + *six6* additive+ *vgll3* parental + *six6* mother + condition.mother | 9 | 5024.243 |
| SMR + *vgll3* additive + *six6* additive+ *vgll3* parental + length.mother | 8 | 5030.794 |
| SMR + *vgll3* additive + *six6* additive+ *vgll3* parental + weight.mother | 8 | 5031.012 |
| SMR + *vgll3* additive + *six6* additive+ *vgll3* parental + condition.mother | 8 | 5029.418 |

**Supplementary table 6:** Linear models testing the effect of maternal genotype on maternal size variables (length and weight) and condition.

| | Covariates | Estimate | Std. Error | t value | Pr(>\|t\|) |
|---|---|---|---|---|---|
| Mother length | (Intercept) | 575.218 | 3.138 | 183.324 | |
| | *vgll3* maternal | 19.190 | 8.821 | 2.176 | 0.031 |
| | *six6* maternal | 20.921 | 7.344 | 2.849 | 0.005 |
| Mother weight | (Intercept) | 1.361 | 0.022 | 61.548 | |
| | *vgll3* maternal | 0.101 | 0.062 | 1.618 | 0.108 |
| | *six6* maternal | 0.195 | 0.052 | 3.759 | <0.001 |
| Mother condition | (Intercept) | 5.908 | 0.035 | 167.039 | |
| | *vgll3* maternal | -0.108 | 0.099 | -1.086 | 0.279 |
| | *six6* maternal | 0.245 | 0.083 | 2.956 | 0.004 |
